# A multi-scale model reveals cellular and physiological mechanisms underlying hyperpolarisation-gated synaptic plasticity

**DOI:** 10.1101/418228

**Authors:** Yubin Xie, Marcel Kazmierczyk, Bruce P. Graham, Mayank B. Dutia, Melanie I. Stefan, Mayank B. Dutia

## Abstract

Neurons in the medial vestibular nucleus (MVN) display hyperpolarisation-gated synaptic plasticity, where inhibition believed to come from cerebellar cortical Purkinje cells can induce long-term potentiation (LTP) or long-term depression (LTD) of vestibular nerve afferent synapses. This phenomenon is thought to underlie the plasticity of the vestibulo-ocular reflex (VOR). The molecular and cellular mechanisms involved are largely unknown. Here we present a novel multi-scale computational model, which captures both electrophysiological and biochemical signalling at vestibular nerve synapses on proximal dendrites of the MVN neuron. We show that AMPA receptor phosphorylation at the vestibular synapse depends in complex ways on dendritic calcium influx, which is in turn shaped by patterns of post-synaptic hyperpolarisation and vestibular nerve stimulation. Hyperpolarisation-gated synaptic plasticity critically depends on the activation of LVA calcium channels and on the interplay between CaMKII and PP2B in dendrites of the post-synaptic MVN cell. The extent and direction of synaptic plasticity depend on the strength and duration of hyperpolarisation, and on the relative timing of hyperpolarisation and vestibular nerve stimulation. The multi-scale model thus enables us to explore in detail the interactions between electrophysiological activation and post-synaptic biochemical reaction systems. More generally, this model has the potential to address a wide range of questions about neural signal integration, post-synaptic biochemical reaction systems and plasticity.

## 1 Introduction

Inhibition in neural circuits plays a fundamental role in modulating the activity and dynamic responsiveness of neurons [Brock et al., 1952, Isaacson and Scanziani, 2011, Gidon and Segev, 2012, Bloss et al., 2016, Doron et al., 2017, Hull, 2017]. In cerebellum-dependent motor learning, inhibitory projections of Purkinje neurons in the cerebellar cortex to neurons in the vestibular nucleus (VN) and deep cerebellar nucleus (DCN), are essential for the induction of adaptive plasticity in extra-cerebellar motor pathways. Purkinje neurons integrate afferent signals from a wide range of sensory systems and inputs from inferior olive neurons, which encode ‘errors’ in ongoing motor acts [Ito, 2002, Yakusheva et al., 2007, Wulff et al., 2009]. While initial models proposed that motor memories were acquired and retained within the cerebellar cortex [Ito, 1982], much experimental and theoretical evidence now shows that motor learning involves the transfer of cortically-acquired motor memories to sub-cortical and extra-cerebellar structures for consolidation and retention [Nagao and Kitazawa, 2003, Kassardjian et al., 2005, Shutoh et al., 2006, Porrill and Dean, 2007, Anzai et al., 2010, Menzies et al., 2010, Clopath et al., 2014, Carcaud et al., 2017]. Many studies have focussed on the neural circuitry underlying the horizontal vestibulo-ocular reflex (VOR) in particular. The VOR is the reflex by which the eyes move in the opposite direction to the head in order to maintain a stable image. The VOR is a robust and tractable experimental system in which to study cerebellum-dependent motor learning [Blazquez et al., 2004, Boyden et al., 2004, du Lac et al., 1995, Miller et al., 2005, Broussard et al., 2011, Shin et al., 2014]. Errors in the gain of the VOR, where the evoked eye movements are larger or smaller than those required to precisely compensate for head movements, result in a slippage of the optical image on the retina and the loss of visual field stability. In response to ‘retinal slip’ errors encoded by inferior olive neurons, Purkinje cells in the cerebellar flocculus are believed to induce potentiation or depression of the vestibular nerve afferent synapses on medial VN (MVN) neurons, so increasing or decreasing the gain of the VOR appropriately to cancel retinal slip [Raymond and Lisberger, 1998, Medina, 2010, Clopath et al., 2014, Carcaud et al., 2017].

Recent studies have revealed a distinct form of hyperpolarisation-gated synaptic plasticity in MVN and DCN neurons, where inhibition presumed to be mediated by Purkinje cell synapses, induces long-term potentiation (LTP) or depression (LTD) of heterologous excitatory, glutamatergic synapses on these neurons [Pugh and Raman, 2009, McElvain et al., 2010]. This represents therefore a plausible cellular mechanism by which cerebellar Purkinje cells may induce lasting changes in sub-cortical neural networks during the consolidation of a learned motor memory [Kassardjian et al., 2005, Shutoh et al., 2006, Anzai et al., 2010].

The cellular and molecular mechanisms that mediate hyperpolarisation-gated plasticity are largely unknown. In both DCN and MVN neurons, excitatory synaptic stimulation paired with a pattern of hyperpolarisation and release from hyperpolarisation, has been shown to induce LTP in the excitatory synapses [Pugh and Raman, 2009, McElvain et al., 2010]. In DCN neurons the activation of low-threshold T type calcium channels (LVCa channels) upon the release of hyperpolarisation, is required for LTP [Person and Raman, 2010]. T type calcium channels are also expressed in MVN neurons [Serafin et al., 1991a,b, Him and Dutia, 2001, Engbers et al., 2013], and interestingly these channels are significantly up-regulated during vestibular compensation, the behavioural recovery that takes place after vestibular deafferentation [Him and Dutia, 2001, Straka et al., 2005, Menzies et al., 2010].

Vestibular nerve and Purkinje cell synapses have recently been demonstrated to be closely apposed on dendrites of parvocellular MVN neurons, providing the anatomical substrate for a close spatial interaction between convergent inhibitory and excitatory synapses in these neurons [Matsuno et al., 2016]. To investigate the cellular and molecular mechanisms that might underlie such interactions, we developed a novel multi-scale model. This model integrates the electrophysiological model of a Type B MVN neuron developed by Quadroni and Knopfel [1994] with biochemical models of postsynaptic calcium signalling and the subsequent activation of LTP- or LTD-inducing pathways, originally designed to model plasticity in hippocampal dendritic spines [Stefan et al., 2008, Li et al., 2012, Mattioni and Le Novère, 2013]. The resulting multi-scale model allowed us to explore the molecular mechanisms invoked by the patterns of inhibition and excitatory stimulation that mediate hyperpolarisation-gated synaptic plasticity. We show that different types of synaptic activity are associated with different sources of calcium influx, with low-voltage activated calcium channels (LVACCs) becoming the dominant route of calcium entry during hyperpolarisation-gated synaptic plasticity. We also show that AMPA receptor phosphorylation (which we use as a readout for synaptic potentiation) shows a highly sigmoidal response to calcium, and that this response is mainly determined by the balance between kinase and phosphatase activity. We further show that the extent and direction of hyperpolarisation-gated synaptic plasticity depend on the strength and duration of the hyperpolarising stimulus, as well as on the relative timing between hyperpolarisation and excitation. Our multiscale model provides a novel in-silico test bed for further investigations into the interplay between inhibition, excitation and biochemical processes in synaptic plasticity.

## 2 Results

### 2.1 Biochemical Pathway Model of Synaptic Plasticity

We modelled the biochemical pathways underlying synaptic plasticity based on an earlier model by Li et al. [2012] (see Methods for details). A Systems Biology Graphical Notation (SBGN) diagram [Le Novère et al., 2009] describing the main components of our model is shown in Fig. 1B. Calcium enters through voltage-dependent calcium channels (VDCCs) and NMDA receptors (shown in grey, not explicitly included in the chemical model). Calcium then binds to and activates calmodulin (shown in yellow). Active (R-state) calmodulin can then activate either the calmodulin-dependent protein kinase II (CaMKII) pathway (shown in red) or the Protein phosphatase 2B (PP2B) (also known as calcineurin (CaN)) pathway (shown in blue). CaMKII is active when it is bound to calmodulin or autophosphorylated. Active CaMKII phosphorylates AMPA receptors, leading to enhanced AMPAR activity. In the PP2B pathway, calmodulin-bound PP2B dephosphorylates DARPP-32, and thereby releases PP1 from inhibition. Active PP1 can dephosphorylate AMPA receptors, counteracting the effect of CaMKII. PP1 also directly dephosphorylates CaMKII itself. The PKA pathway (shown in pink) can reduce dephosphorylation of AMPA receptors by activating DARPP-32. Taken together, the model captures the sequence of events leading from calcium influx to changes in AMPAR phosphorylation state.

**Figure 1:**
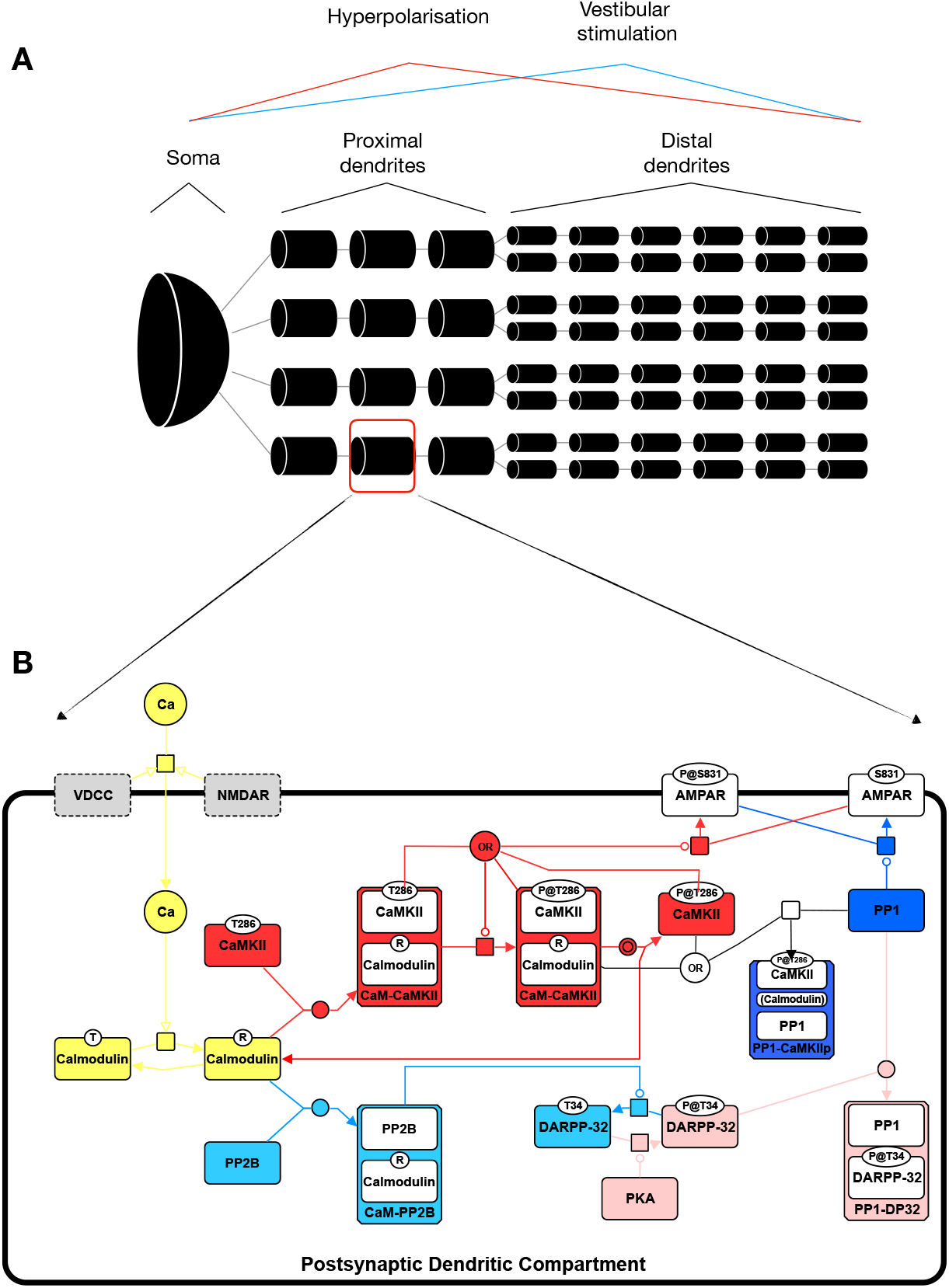
Simplified schematic of the models used in this work. **A**. NEURON model with 61 electrical compartments, representing the soma, proximal dendrites, and distal dendrites. **B**. Chemical model with the key molecular species and reactions that underlie calcium-dependent synaptic plasticity.

### 2.2 AMPA receptor phosphorylation shows a complex dependency on calcium concentration

In order to establish the usefulness of our biochemical model as a model of synaptic plasticity, we first examined the dependence of AMPA receptor phosphorylation on calcium concentration. To this end, we determined the effects of a step change in [Ca^2+^] from baseline at equilibrium to a range of concentrations over the range 4 × 10^−10^ M to 4 × 10^−7^ M (Fig. 2). The time course of changes in the active molecular species in the system were observed over 1600 s after the step change in [Ca^2+^]. As output, we computed the AMPAR phosphorylation ratio (phosphorylated AMPAR over total AMPAR).

**Figure 2:**
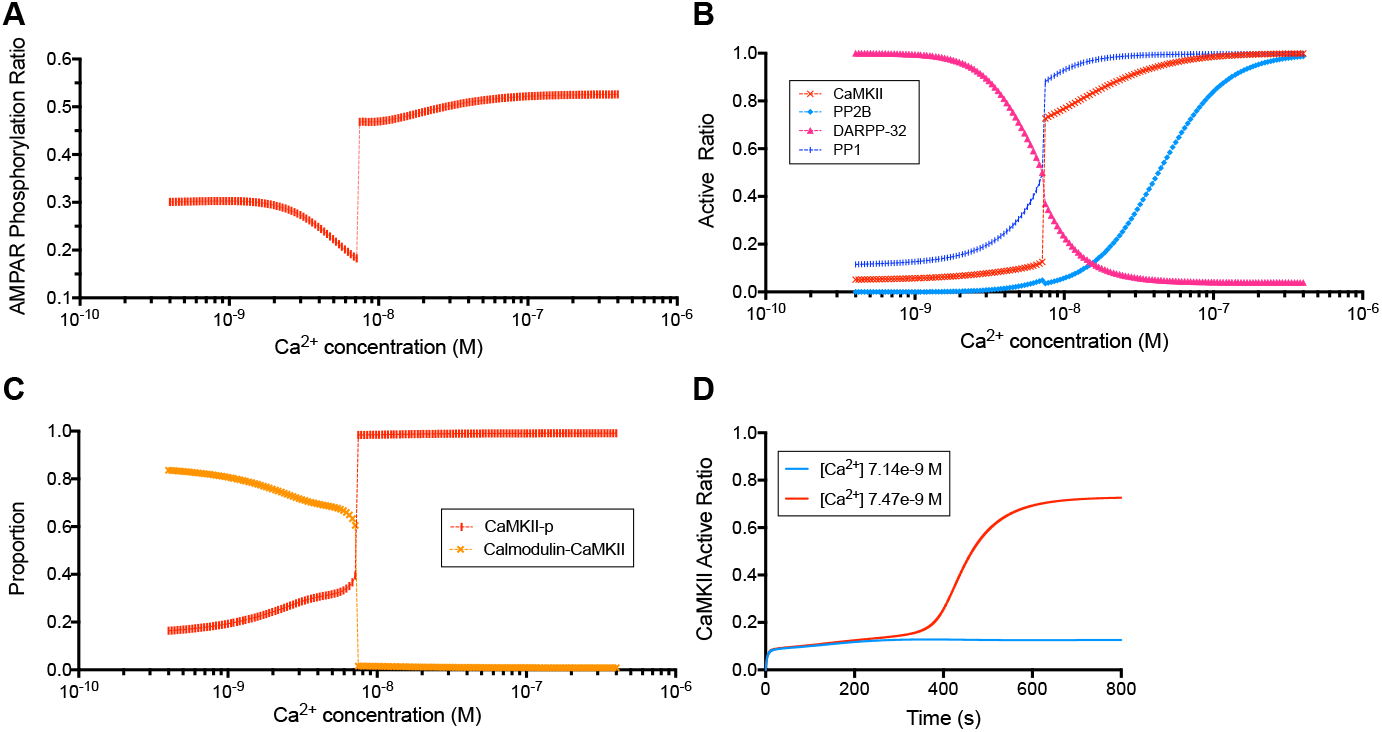
Dependence of AMPAR phosphorylation and relative activation ratios of key molecular species on [Ca^2+^]. **A**. AMPAR phosphorylation ratio (phosphorylated AMPAR over total AMPAR) as a function of [Ca^2+^] over the range 4 × 10^−10^ M to 4 × 10^−7^ M. For each data point the biochemical model was reinitialised and after reaching equilibrium the intracellular [Ca^2+^] was stepped up to successively higher values (logarithmic scan of [Ca^2+^] range, 151 simulations). Each simulation was run for 1600 s, to steady state. Note the initial dephosphorylation of AMPA receptors with increasing [Ca^2+^], which is followed by a switch to a large sustained increase in AMPAR phosphorylation when [Ca^2+^] is increased above a threshold value for inducing long-term potentiation. **B**. Relative activation ratios (concentration of active molecules divided by total molecules) for CaMKII, PP2B, DARPP-32, and PP1 as a function of [Ca^2+^]. **C**. The proportion of the two active CaMKII forms: CaMKII-p (phosphorylated CaMKII) and calmodulin-bound (non-phosphorylated) CaMKII as a function of [Ca^2+^]. D. Time-course of activation of CaMKII over a period of 800 s, at two values of [Ca^2+^] on either side of the threshold concentration.

As can be seen from Fig. 2A, the relationship between [Ca^2+^] and AMPAR phosphorylation is complex. At very low calcium concentrations there is a baseline level of AMPAR phosphorylation of about 30 %. Moderate increases in calcium concentrations (to the nanomolar range) lead to a decrease in AMPAR phosphorylation ratio. At [Ca^2+^] of between 7 and 7.5 × 10^−9^ M however, there is a sharp increase in AMPAR phosphorylation ratio, so that at [Ca^2+^] values of 7.5 × 10^−9^ M and higher about 50% of total AMPA receptors are phosphorylated at the end of the 1600 s simulation period.

In order to identify the molecular determinants of the changes in AMPA receptor phosphorylation, we examined the activation ratios (active over total concentrations) of other key molecular species in the system over the same range of Ca^2+^ concentrations (Fig. 2B). The activation patterns of kinases and phos-phatases show sigmoidal responses to initial calcium concentration, with different directions, incline, and transition concentrations. At sub-threshold Ca^2+^ concentrations, the dominant active form of CaMKII is calmodulin-bound (non-phosphorylated) CaMKII, whereas at higher Ca^2+^ phosphorylation becomes the main contributor to CaMKII activity (Fig. 2C). In order to highlight the bistable response pattern of CaMKII, we ran time courses at calcium concentrations just below and just above the threshold concentration at which the system switches (Fig. 2D). The bi-stable response is also present when calmodulin concentration is reduced, although the sharpness of the response and the calcium concentration at which the system changes between states vary (Fig. S12).

The same bistable pattern of kinase and phosphatase behaviour has been found in the earlier models that this model builds on Stefan et al. [2008], Li et al. [2012]. Thus, our model reproduces earlier findings, and extends them to describe AMPAR phosphorylation in response to Calcium signalling. Since dephosphorylation of AMPA receptors is associated with LTD and AMPA receptor phosphorylation with LTP, this behaviour of the biochemical system is consistent with the widely accepted model by which a moderate increase in [Ca^2+^] will result in the depression of excitatory synapses, and a greater increase will result in their potentiation [Lisman, 1989, Malenka, 1994]. We therefore used AMPA receptor phosphorylation ratio as a readout in our subsequent simulation experiments.

We next implemented the biochemical model within dendritic compartments of the MVN neuron, in order to study LTD and LTP of the excitatory vestibular nerve synapses that impinge upon the MVN neuron.

### 2.3 Dendritic [Ca^2+^] profiles induced by vestibular synaptic stimulation with and without hyperpolarisation

We next studied how different stimulation protocols affect calcium dynamics in the proximal dendrites of the MVN neuron. For this, we used a multi-compartmental electrophysiological model of an MVN Type B neuron [Quadroni and Knopfel, 1994, Graham et al., 2009] in the NEURON software [Carnevale and Hines, 2006] (Fig 1A). We stimulated each neuronal compartment in the model by either activating the excitatory vestibular synapses (Vestibular Stimulation protocol, “VS”) or by activating the excitatory vestibular synapses together with hyperpolarisation of the cell membrane (Hyperpolarisation + Vestibular Stimulation protocol, “H+VS”).

The results are shown in Fig. 3. The MVN neuron had a resting firing rate of around 24 Hz before vestibular stimulation, similar to the resting activity observed in experimental studies [Johnston et al., 1994, McElvain et al., 2010] (Fig. 3 A). Stimulation of the vestibular nerve input at 100Hz (“VS protocol”) evoked a firing rate of around 56 Hz, similar to the response evoked by vestibular nerve stimulation in the study of McElvain et al. [2010]. There was a significant increase in [Ca^2+^] for the duration of the vestibular nerve stimulation, with the result that the total dendritic [Ca^2+^] was elevated to 2.3 times the baseline level (Fig. 3 B, D). Analysis of the individual dendritic ion channel currents (Fig. 3C) showed that at rest, the Ca^2+^ influx was mediated predominantly by LVA Ca^2+^ channels and by HVA Ca^2+^ channels that were activated during each action potential. By contrast during VS stimulation, the activation of synaptic NMDA channels in addition to the LVA and HVA channels, led to a significant increase of dendritic Ca^2+^ influx for the duration of the vestibular stimulation (Fig. 3C, D).

**Figure 3:**
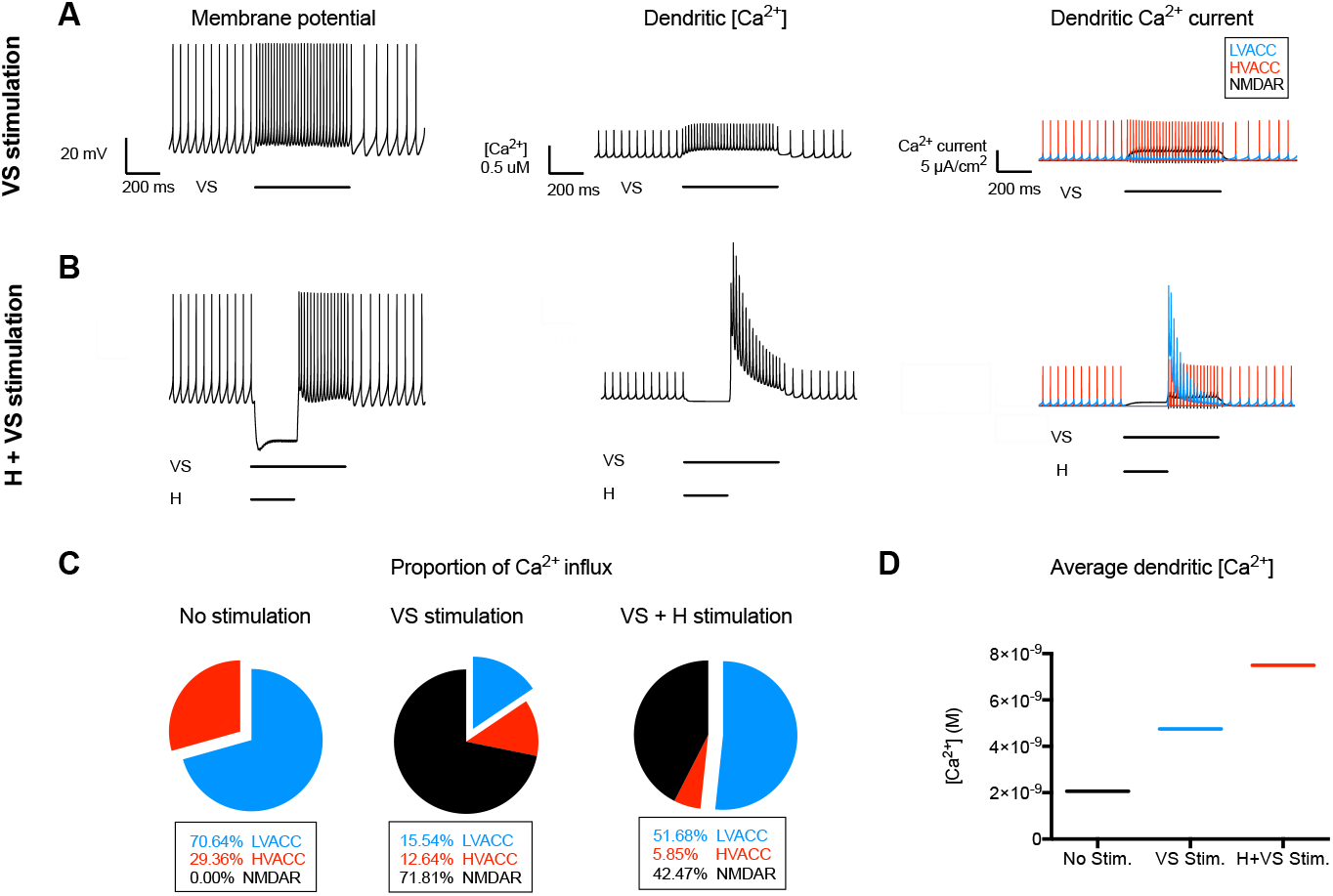
Effects of VS and H+VS stimulus protocols on membrane potential and [Ca^2+^] in a dendritic compartment of a Type B MVN cell. The membrane potential and action potential firing (left panel), total [Ca^2+^] profiles (middle panel) and Ca^2+^ currents dynamics (right panel) in a single dendritic compartment of a NEURON model of a Type B MVN cell in response to vestibular nerve synapse stimulation at 100 Hz for 550 ms (VS protocol, **A**), and to vestibular stimulation combined with membrane hyperpolarisation for the first 250 ms of the stimulation period (H+VS protocol, **B**). **C**. the contribution of low-voltage activated calcium channels (LVACC), high-voltage activated calcium channels (HVACC) and NMDA receptor channels (NMDAR) to the changes in [Ca^2+^] observed in the two stimulus protocols. Note the activation of HVACC during action potential firing, and the activation of synaptic NMDAR during VS stimulation. During H+VS stimulation, the release of the membrane hyperpolarisation results in a large activation of LVACC Ca^2+^ currents. **D**. Total dendritic [Ca^2+^] averaged over the simulation period for the VS and H+VS stimulus protocols after scaling due to Ca^2+^ buffering effect (Fig. 11).

As shown in Fig. 3B, in response to the H+VS stimulation protocol there was an initial hyperpolarisation of around 25 mV over the first 250 ms, which was followed by a rebound depolarisation and increase in firing rate upon the release of inhibition. Dendritic [Ca^2+^] decreased initially during the period of inhibitory stimulation, but the large influx of [Ca^2+^] during the rebound depolarisation resulted in the net dendritic [Ca^2+^] being elevated to a level 3.64 times the baseline (Fig. 3D). The Ca^2+^ influx from LVA channels activated upon the release of inhibition was the predominant source of dendritic Ca^2+^ during the H+VS stimulation. Together with the NMDA receptor mediated influx, this led to the highly elevated levels of dendritic [Ca^2+^] (Fig. 3D).

Together, these results demonstrate that during vestibular nerve stimulation alone (VS protocol) the activation of synaptic NMDA receptors causes a moderate elevation of dendritic [Ca^2+^], while pairing hyperpolarisation with vestibular nerve stimulation in the H+VS protocol led to the elevation of dendritic [Ca^2+^] to a substantially higher level. Thus, postsynaptic hyperpolarisation has a marked effect on dendritic [Ca^2+^] influx, when the activation of LVA channels coincides with activation of the synaptic NMDA receptors.

### 2.4 Multi-scale modelling reveals molecular pathways activated in response to vestibular synaptic stimulation with and without hyperpolarisation

We next examined the effect of calcium influx under the VS and H+VS protocols on downstream signalling pathways. In order to do this, we combined the NEURON and COPASI models described above into one multi-scale modelling pipeline. Specifically, we first used the NEURON model to simulate the VS and H+VS protocols, as described above. The readout from the neuron model was a profile of intracellular calcium concentration over time. These profiles were then used as an input for a COPASI simulation of the calcium-dependent biochemical signalling pathways as described earlier. The interface between NEURON and COPASI was coded in Python and included two steps to facilitate a smooth conversion from electrical to biochemical model. First, we binned time steps from the NEURON model into larger time intervals, because biochemical simulations happen at a larger time scale and therefore do not require the same temporal resolution as electrophysiological models. Second, we applied a scaling factor in order to account for the fact that a portion of the calcium entering the cell will immediately bind to intracellular calcium buffers not explicitly represented in our biochemical model, and the effective calcium concentration available for calmodulin binding will therefore be smaller than the overall calcium concentration.

The outcomes of the multi-scale simulations are shown in Fig. 4. The dendritic [Ca^2+^] profiles induced by the VS and H+VS stimulation protocols had substantially different effects on the key components of the biochemical model and the resultant phosphorylation of AMPA receptors. In particular the moderate rise of dendritic [Ca^2+^] induced by the VS stimulation led to a modest activation of CaMKII and PP2B (Fig. 4 A), and a sustained decrease in the level of AMPA receptor phosphorylation over the course of the simulation (Fig. 4C). By contrast, the greater rise in [Ca^2+^] induced by the H+VS stimulation led to a marked activation of PP2B and CaMKII (Fig. 4B). Following an initial decrease in active AMPA receptor phosphorylation ratio, the H+VS stimulation protocol induced a large sustained increase in AMPA receptor phosphorylation level (Fig. 4D). Presumably the rise in [Ca^2+^] evoked by the H+VS protocol was greater than that required to trigger auto-phosphorylation of CaMKII and the consequent high level of phosphorylation of the AMPA receptors, while in the VS protocol the modest rise in [Ca^2+^] was insufficient to trigger this ‘‘switchlike” behaviour. The VS and H+VS protocols thus had opposite effects on synaptic AMPA receptors. This is consistent with the experimental observation of synaptic LTD and LTP in the study of McElvain et al. [2010].

**Figure 4:**
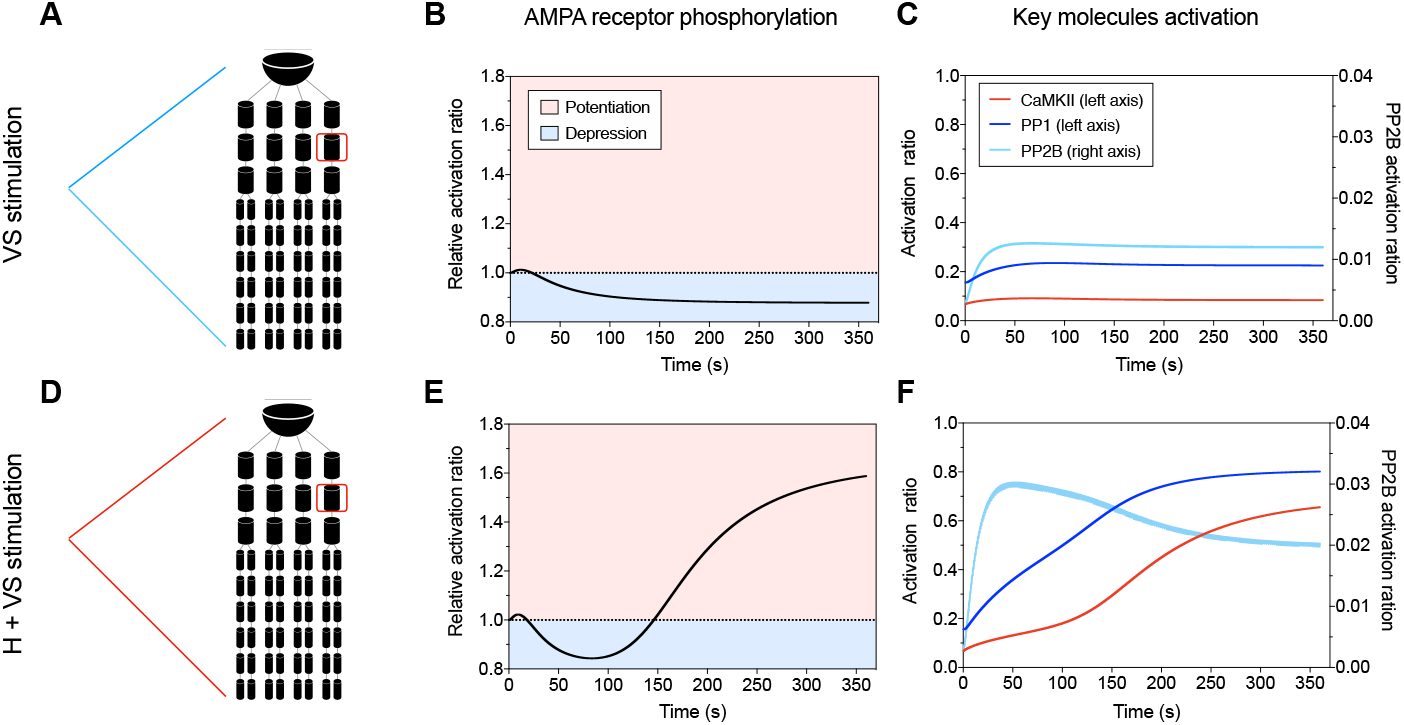
Time-course of activation of key molecular species and AMPAR phosphorylation in response to VS and H+VS stimulation. Left part is the changes in active ratios of CaMKII, PP1 and PP2B in response to the VS stimulation protocol (**A**) and H+VS stimulation (**B**). Right part corresponding changes in AMPAR phosphorylation in response to the VS stimulation protocol (**A**) and H+VS stimulation protocol (**B**). Note that the VS protocol evokes a moderate activation of the key molecular species and results in a net de-phosphorylation of AMPAR over the period of stimulation, while the H+VS protocol induces a marked activation of PP1 and CaMKII, and results in a sustained phosphorylation of AMPA receptors.

Taken together, vestibular stimulation following hyperpolarisation results in markedly increased calcium influx, activation of calcium-dependent kinases over phosphatases and a resulting net increase in AMPA receptor phosphorylation. In contrast, vestibular stimulation alone results in a more moderate calcium influx, a different balance of kinase and phosphatase activity, and a net dephosphorylation of AMPA receptors.

### 2.5 LVA channel density determines dendritic calcium dynamics and hyperpolarisation-gated synaptic plasticity

To examine further the role of MVN LVA channels in mediating hyperpolarisation-gated plasticity, we explored the relationship between LVA channel density, the post-hyperpolarisation rebound calcium influx and effects on synaptic AMPAR phosphorylation. We compared the effects of the VS and H+VS protocols on AMPAR phosphorylation ratio in a “Type B_high_LVA_ neuron” model, in which the dendritic LVA channel density is significantly higher than the normal Type B neuron [Quadroni and Knopfel, 1994] (Table 2), and in the hypothetical “Type B_low_LVA_ neuron” in which the LVA channel density was reduced to 50% of that in the normal Type B neuron (Table 2). As expected, in the Type B_high LV_A neuron the post-hyperpolarisation rebound firing and the dendritic Ca^2+^ influx were significantly greater than in the normal Type B neuron, reflecting the higher density of dendritic LVA channels in this cell type. By contrast the Type Bι_ow LV_A neuron showed a much smaller rebound depolarisation and reduced dendritic Ca^2+^ influx than the normal Type B neuron (Fig. 5). As shown in Fig. 6, the effects of the VS and H+VS protocols on AMPAR phosphorylation levels in these neuron types were also markedly different. In the Type B_high LV_A neuron, the VS stimulation protocol caused a slightly more pronounced decline in AMPA receptor phosphorylation level than in the “normal” Type B neuron. The H+VS stimulation protocol evoked a larger, more rapid increase in AMPA receptor phosphorylation (Fig. 6). By contrast in the Type B_low_LVA_ neuron VS stimulation caused a slower, less marked decrease in AMPAR phosphorylation levels than in the normal Type B neuron, while the H+VS stimulation did not result in increased AMPAR phosphorylation, instead producing a slight decline.

**Figure 5:**
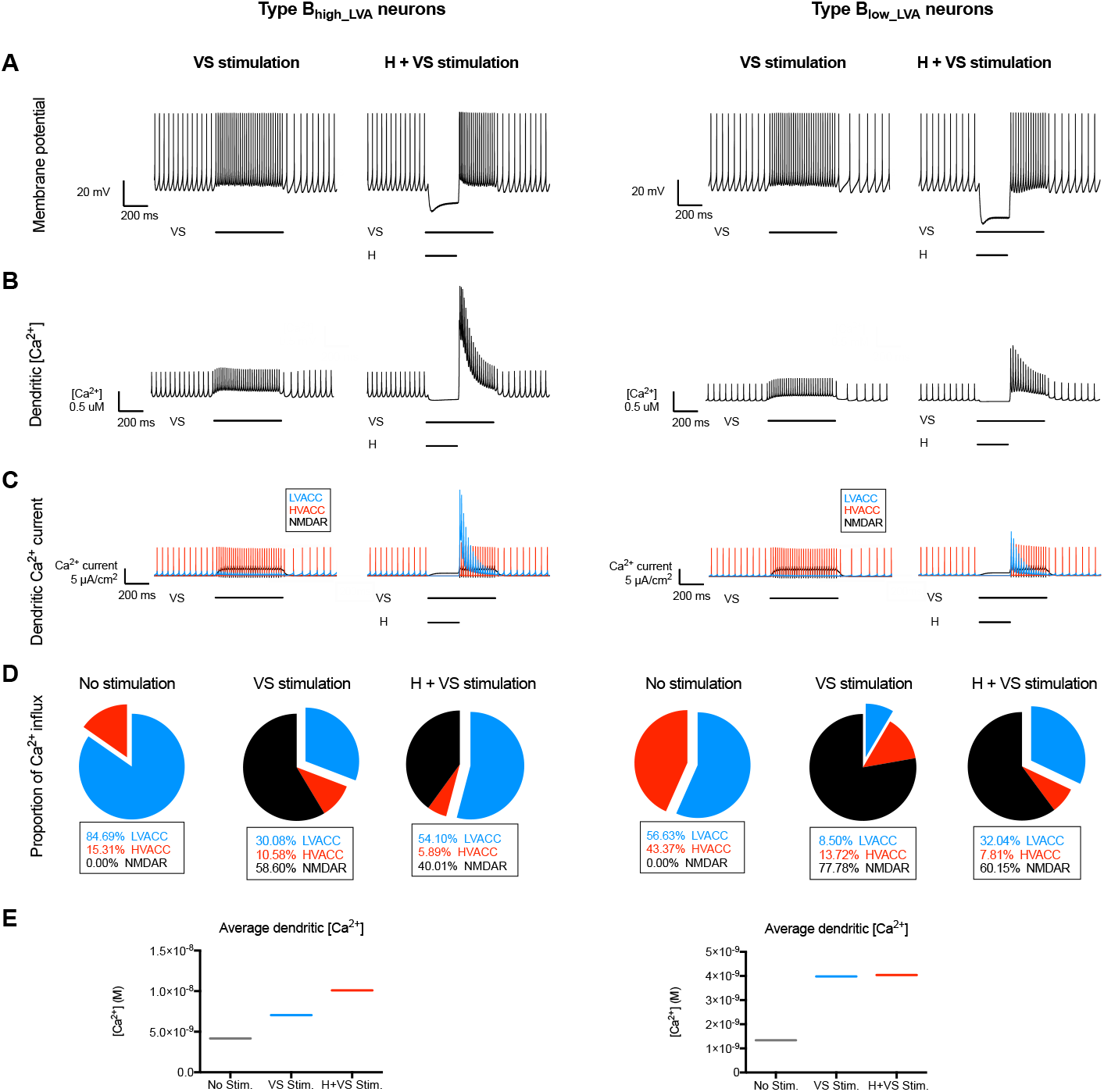
Effects of VS and H+VS stimulus protocols on membrane potential and [Ca^2+^] in a dendritic compartment of a Type B_high_LVA_ or Type B_low_LVA_ MVN cell. The same stimulation protocols and analysis as in Fig. 3 are applied here. In the Type B_high_LV_A neuron the post-hyperpolarisation rebound firing and the dendritic Ca^2+^ influx were significantly greater than in the normal Type B neuron, while the Type B_low_LVA_ neuron showed a much smaller rebound depolarisation and reduced dendritic Ca^2+^ influx than the normal Type B neuron. Note that because the types of neurons differ in LVA density, they have different basal calcium concentrations, and a direct comparison of absolute values between types of neurons is therefore difficult.

**Figure 6:**
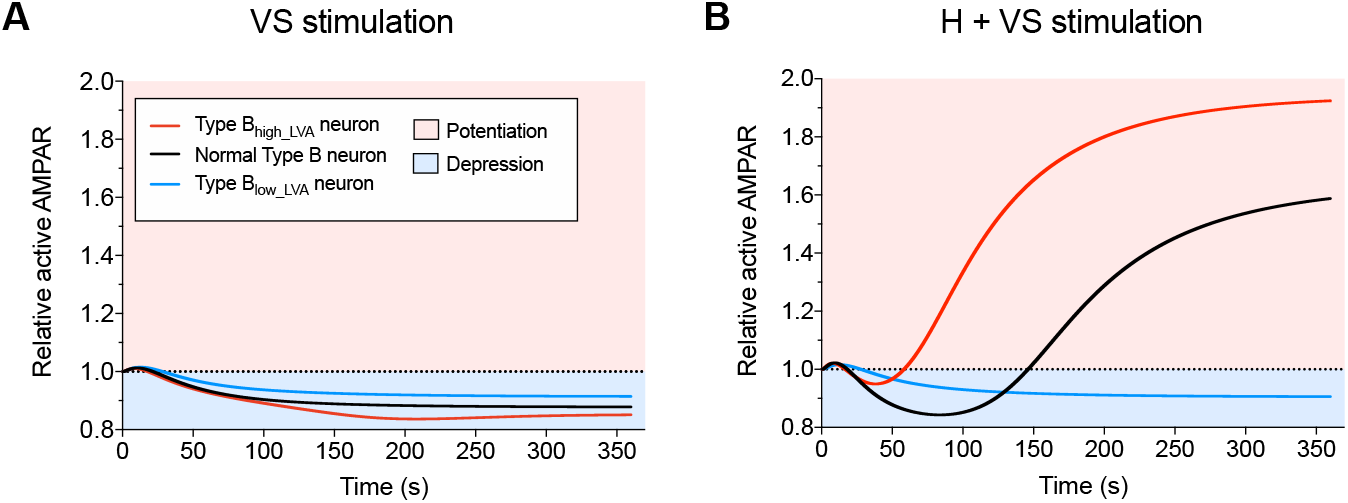
Effects of VS and H+VS stimulation protocols on AMPAR phosphorylation depend on LVA channel expression in MVN neurons. Effects of the VS (**A**) and H+VS (**B**) stimulation protocols on AMPAR phosphorylation in a normal Type B MVN neuron, a Type B neuron expressing a high level of LVACC (Type B _high_LVA_ cell), and a hypothetical class of MVN neuron with low expression of LVACC (Type B_low_LVA_ cell).

Together, these results indicate that the density of LVA channels is a major determinant of dendritic calcium dynamics and hyperpolarisation-gated synaptic plasticity.

### 2.6 Hyperpolarisation-gated synaptic plasticity depends on the balance of kinases and phosphatases

Given the importance of dendritic calcium influx and its biochemical effects in regulating AMPA receptor phosphorylation levels, we next examined the effects of varying the relative concentrations of the two key molecules CaMKII and PP2B, on the response of the normal MVN Type B neuron to the H+VS stimulation protocol (Fig. 7). We ran a total of 9 simulations where we reduced the intracellular concentrations of CaMKII and PP2B to 66% and 33% of the base model independently and in combination, to explore the effects of changing the absolute levels of these molecules as well as their relative ratios. As shown in Fig. 7, total CaMKII concentration is the main determinant of AMPAR phosphorylation level, although the balance between kinase and phosphatase concentration also plays a role. At normally high CaMKII concentrations (Fig. 7A), there is always an increase in AMPAR phosphorylation, though that increase is more pronounced when PP2B concentrations is reduced. Interestingly, when CaMKII concentration is slightly reduced (Fig. 7B), this can produce either an increase or a decrease in AMPAR phosphorylation, depending on PP2B concentration. At low CaMKII levels (Fig. 7C), the kinase activity is no longer enough to induce a net increase in AMAPR phosphorylation at normal or 66% of normal levels of PP2B. Only if PP2B concentration is also lowered to 33% of its base model concentration can we see an increase in AMPAR phosphorylation, but this increase is slow and moderate in scale.

**Figure 7:**
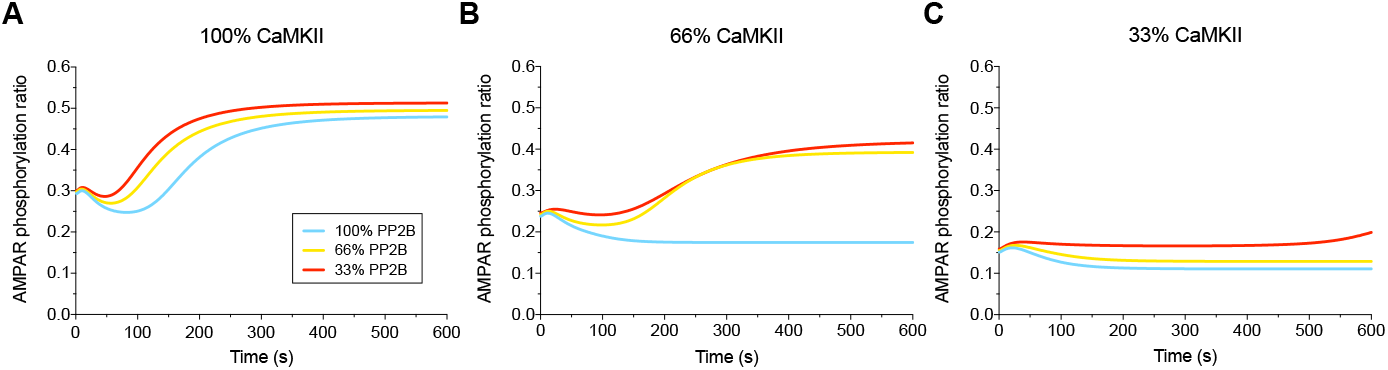
Dependence of AMPAR phosphorylation response on CaMKII, PP1 and PP2B levels. Effects of varying the concentrations of CaMKII and PP2B in a normal Type B MVN neuron on the phosphorylation of AMPAR in response to the H+VS stimulation protocol. Simulations were run where CaMKII and PP2B concentrations were set to 100%, 66% and 33% of those typically found in a neuron.

These findings demonstrate the importance of CaMKII as a key molecular species in mediating hyperpolarisation-gated plasticity, and show that the net effectiveness of the H+VS stimulation protocol is critically dependent on the balance between the kinase and its antagonist phosphatase PP2B.

### 2.7 Hyperpolarisation-gated plasticity depends on the strength, duration, and relative timing of the hyperpolarising stimulus

Finally we explored what features of the hyperpolarising stimulus are important for determining the strength and direction of synaptic plasticity. We investigated this by simulating repeated combinations of inhibitory stimuli for 600 s each. Across simulations, we varied either the strength, duration, or relative timing of the hyperpolarising stimulus (corresponding to 600, 600 and 400 inhibition/excitation pairs in 600s), while keeping the other two parameters and the parameters governing the excitatory stimulus constant.

We first varied the amplitude of the hyperpolarising stimulus. In order to do this, we used our multi-scale model to simulate a H+VS protocol in normal Type B MVN neurons with hyperpolarisation strengths ranging from 0 pA to 600pA (Fig. 8A). In all cases, each excitatory pulse lasted for 550ms. Each hyperpolarising stimulus started at the same time as the excitatory stimulus and lasted for 250 ms.

**Figure 8:**
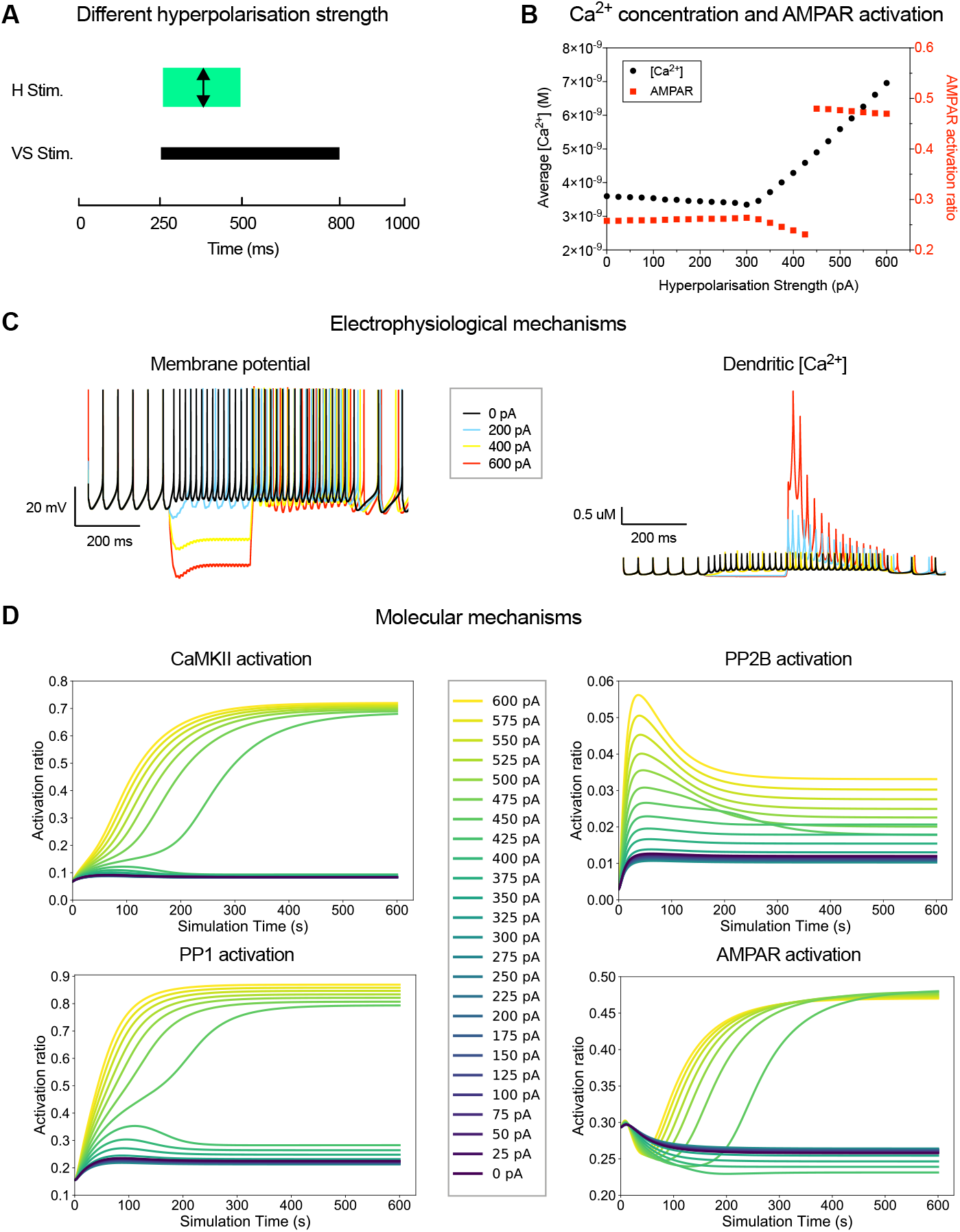
Effects of varying hyperpolarisation strength on AMPA receptor activation. **A**. Simulation set-up: Hyperpolarising stimuli of various strengths were applied to MVN type B neurons. **B**. AMPAR activation at the end of each 600 s simulation plotted as a function of average [Ca^2+^] during the same simulation. **C**. Electrophysiological mechanisms (dynamics of the membrane potential and dendritic [Ca^2+^]) for selected hyperpolarisation strengths. **D**. Molecular changes (activation of kinases and phosphatases and AMPAR phosphorylation over time) for a range of hyperpolarisation strengths.

As shown in Fig. 8B, both dendritic Ca^2+^ concentration and AMPA receptor phosphorylation displayed a complex dependency on the strength of the hyperpolarising stimulus. Small amounts of membrane hyperpolarisation caused essentially no change (or a small degrease) in calcium levels, and no change in AMPA receptor phosphorylation. Intermediate hyperpolarisation strengths resulted in a moderate increase in [Ca^2+^] and a decrease in AMPA receptor phosphorylation. Once hyperpolarisation strength reaches a threshold of 450 pA, the system switches: CaMKII reaches sustained levels of activation, resulting in an overall increase of AMPA receptor phosphorylation. (Fig. 8D).

Next, we varied the duration of the hyperpolarising stimulus. In order to do this, we used our multi-scale model to simulate a H+VS protocol in normal Type B MVN neurons with hyperpolarisation duration ranging from 0 ms to 550 ms (Fig. 9A). In all cases, we used a hyperpolarising stimulus of 475 pA, which started at the same time as a 550 ms excitatory stimulus.

**Figure 9:**
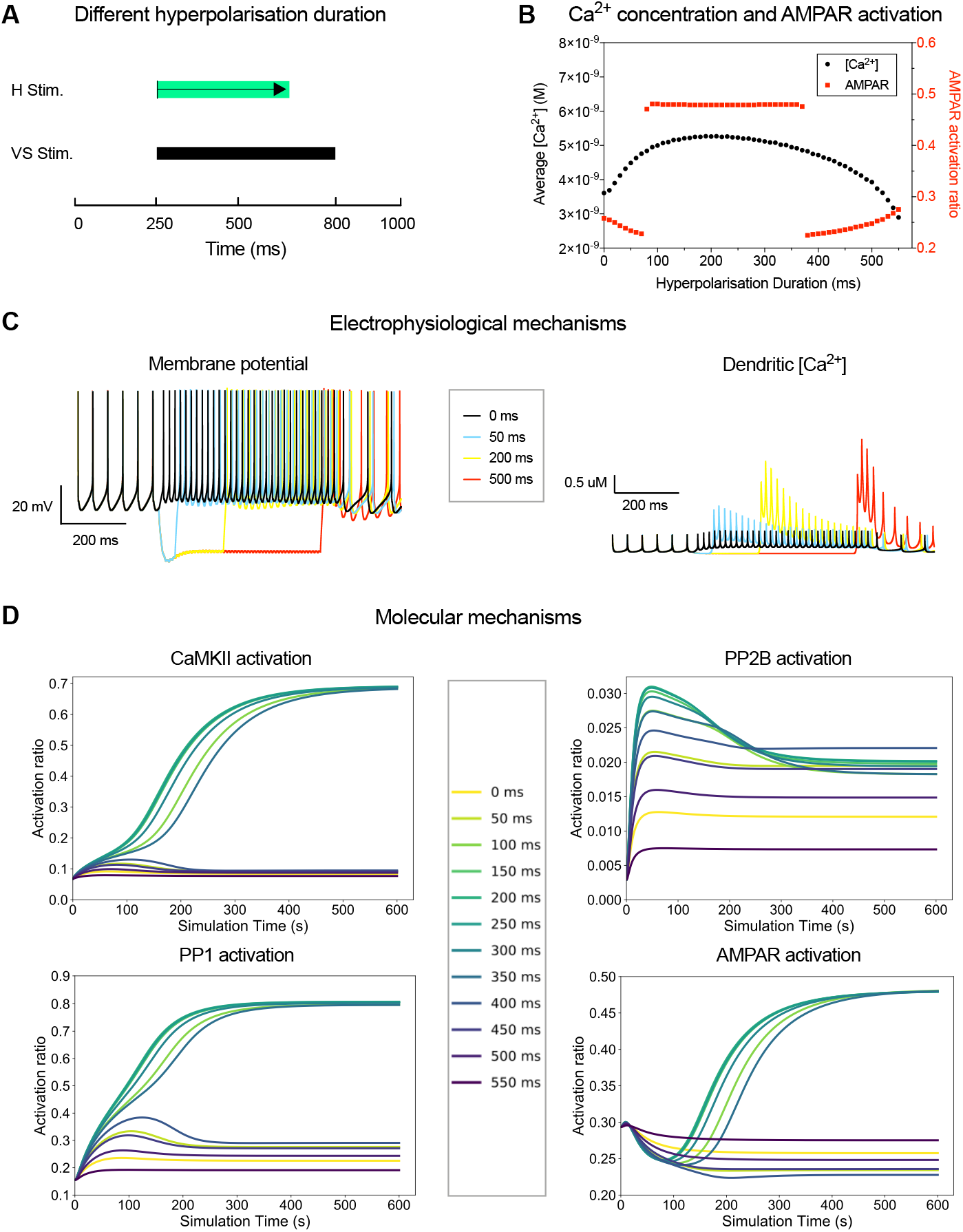
Effects of varying hyperpolarisation duration on AMPA receptor activation. **A**. Simulation set-up: Hyperpolarising stimuli of varying duration were applied to MVN type B neurons. **B**. AMPAR activation at the end of each 600 s simulation plotted as a function of average [Ca^2+^] during the same simulation. **C**. Electrophysiological mechanisms (dynamics of the membrane potential and dendritic [Ca^2+^]) for selected hyperpolarisation durations. D. Molecular changes (activation of kinases and phosphatases and AMPAR phosphorylation over time) for a range of hyperpolarisation durations.

The dendritic [Ca^2+^] response and AMPA receptor phosphorylation showed a complex dependence on the duration of the H stimulus (Fig. 9B). A short period of hyperpolarisation (less than 80 ms) leads to a small increase of [Ca^2+^] in the rebound and no change or a decrease in AMPA receptor phosphorylation. With very long periods of hyperpolarisation that overlap with the excitatory stimulation, the net Ca concentration can even be lower than that without stimulation. This will also lead to a decrease in AMPAR activation. At hyperpolarisation durations in between, there is a robust rise in calcium levels, and a stable “up” state is reached at which almost half of all AMPA receptors are phosphorylated (Fig. 9D).

Finally we varied the timing between hyperpolarising and depolarising stimuli. In order to do this, we used our multi-scale model to simulate a H+VS protocol in normal Type B MVN neurons. We varied the onset of the hyperpolarising stimulus from −500 ms to +250 ms from the onset of the excitatory stimulus (Fig. 10A). In all cases, the amplitude of the hyperpolarising stimulus was 475 pA, and the duration was 250 ms.

**Figure 10:**
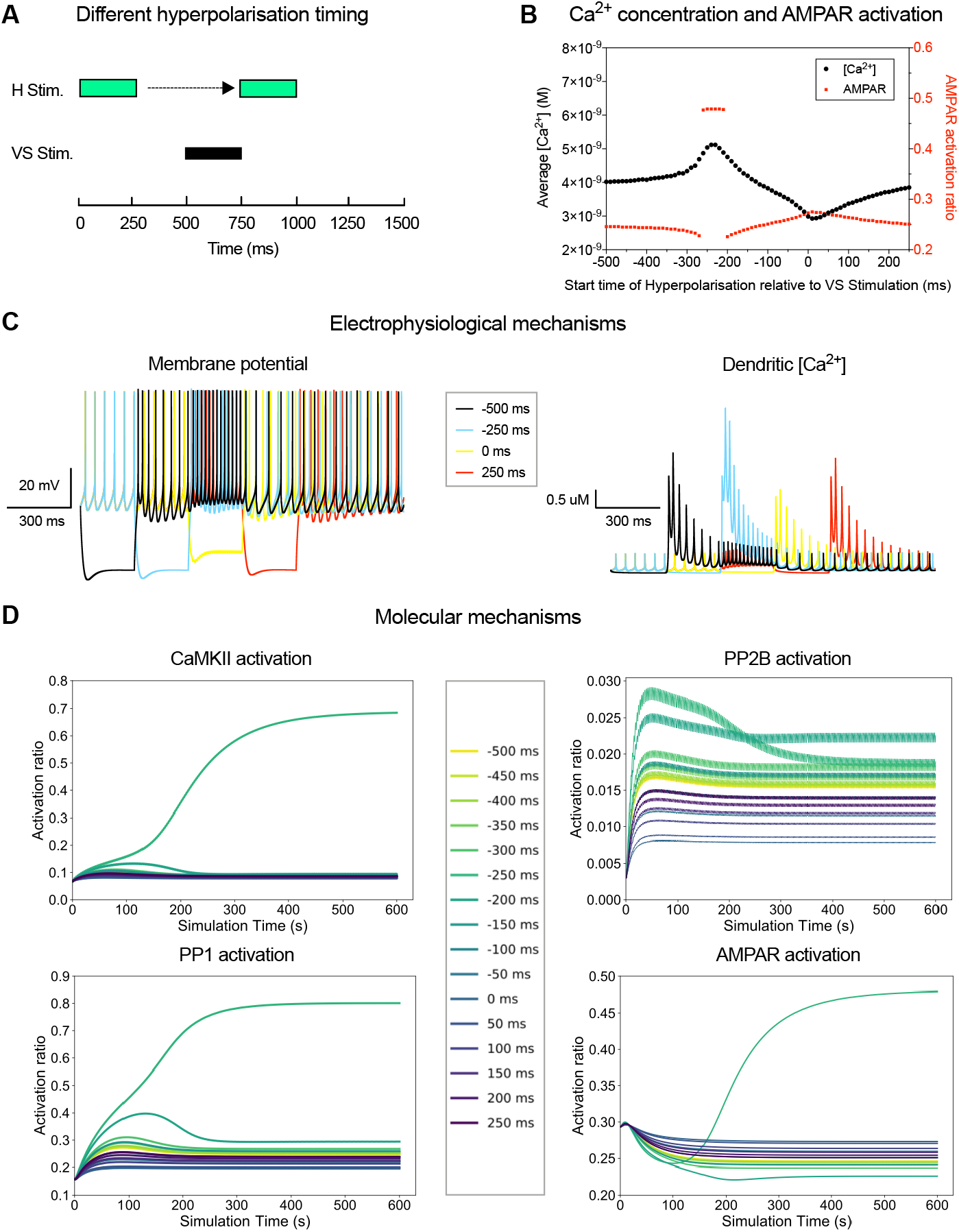
Effects of varying the timing between hyperpolarisation and excitation on AMPA receptor activation. **A**. Simulation set-up: Hyperpolarising and excitatory stimuli were applied at varying relative timing to MVN type **B** neurons. **B**. AMPAR activation at the end of each 600 s simulation plotted as a function of average [Ca^2+^] during the same simulation. **C**. Electrophysiological mechanisms (dynamics of the membrane potential and dendritic [Ca^2+^]) for selected hyperpolarisation onset timings. **D**. Molecular changes (activation of kinases and phosphatases and AMPAR phosphorylation over time) for a range of hyperpolarisation onset timings.

The dendritic [Ca^2+^] response showed a marked dependence on the relative timing of the hyperpolarising and excitatory stimuli (Fig. 10B). A maximal elevation of [Ca^2+^] was seen when the release of the membrane hyperpolarisation corresponded with the start of the VS stimulation. In contrast, dendritic [Ca^2+^] was at its lowest when the onsets of hyperpolarising and excitatory stimuli coincided. Correspondingly, there is a narrow window of relative timings where strong and sustained AMPA receptor phosphorylation is observed (Fig. 10B, D), suggesting that the precise timing of hyperpolarisation and excitation is crucial for hyperpolarisation-gated synaptic plasticity.

### 2.8 Testable predictions

Our model reproduces earlier findings on hyperpolarisation-gated synaptic plasticty McElvain et al. [2010], but in addition generates testable predictions about the underlying mechanism. One prediction is that hyperpolarisation-mediated synaptic plasticity depends crucially on Calcium entry through low-voltage activated calcium channels. We predict that application of selective blockers of those channels (reviewed in Kopecky et al. [2014]) would either abolish the hyperpolarisation-mediated response altogether or skew the balance between LTD and LTP towards LTD.

In addition, we predict that the direction of plasticity is determined by the balance between kinases and phosphatases. It is true that prior work on CaMKII*α* mutant mice (e.g. Silva et al. [1992b,a]) has not described a phenotype relating to plasticity of the vestibulo-ocular reflex. This could be because there is no deficit, or for a number of other reasons: the VOR plasticity phenotype was not explicitly tested for and may be too subtle to be noticed amid the other learning deficits. It is also possible that other mechanisms (biochemical, cellular, or behavioural) compensate for the loss of CaMKII*α*. It would be interesting to repeat the electrophysiological measurements by McElvain et al. McElvain et al. [2010] on slides from CaMKII*α* mutant mice and see whether hyperpolarisation-gated synatpic plasticity is indeed dirsupted on the cellular level. Alternatively, experiments could be performed on wildtype slices after applying specific kinase or phosphatase inhibitors.

Finally, we predict a rather specific non-linear profile of hyperpolarisation-gated synaptic plasticity in response to altering either the strength and duration of hyperpolarisation. Both those response curves are amenable to in-vitro validation.

## 3 Discussion

In this study we developed a detailed multi-scale model incorporating electrophysiological and biochemical processes that regulate AMPA-receptor phosphorylation in MVN neurons, to investigate the interactions between membrane hyperpolarisation and synaptic plasticity. The electrophysiological component of the multi-scale model was based on well-established NEURON models which faithfully reproduce the membrane characteristics of rodent Type B MVN neurons in vitro [Quadroni and Knopfel, 1994]. The biochemical component of the multi-scale model was based on models of calcium signalling and the activation of protein kinases and phosphatases in postsynaptic neuronal compartments [Stefan et al., 2008, Li et al., 2012, Mattioni and Le Novère, 2013]. All those earlier models had been validated against the experimental literature. Although the biochemical models were designed to model hippocampal plasticity, the chemical components are well-conserved between neuron types, and reaction parameters do not change. As a result, our model parameters were well-constrained from the start.

Using our new multi-scale modelling interface, dendritic Ca^2+^ concentration profiles obtained from the NEURON model of the MVN neuron were routed to the biochemical component of the model. This allowed us to model changes in the phosphorylation ratio of AMPA receptors, as a proxy of long-term potentiation or depression of the activated synapses in a dendritic compartment of the MVN neuron.

The multi-scale model thus allowed us to examine for the first time the cellular and molecular mechanisms involved in hyperpolarisation-gated synaptic plasticity, where the synaptic strength of vestibular nerve afferent inputs in MVN neurons is regulated by inhibitory inputs presumably from Purkinje cell synapses [McElvain et al., 2010]. Similar interactions are also seen in DCN neurons [Pugh and Raman, 2009, Person and Raman, 2010], suggesting that hyperpolarisation-gated synaptic plasticity is a fundamental mechanism in cerebellum-dependent motor learning. We used our model to specifically determine the effects on calcium signalling and AMPA receptor regulation induced by experimental protocols that cause either a potentiation or a depression of the vestibular synapses in MVN neurons [McElvain et al., 2010]. During resting activity, dendritic Ca^2+^ concentration showed small transients largely due to Ca^2+^ influx with each action potential through high-threshold Ca^2+^ channels, leading to low baseline levels of AMPA receptor phosphorylation. A train of synaptic stimulation at 100 Hz (VS stimulation protocol) has previously been shown to induce LTD in vestibular synapses [McElvain et al., 2010]. In contrast, if the same train was accompanied by a period of hyperpolarisation (H+VS protocol), then this resulted in LTP [McElvain et al., 2010]. Our model reproduced both effects, with VS stimulation alone causing only moderate increases in calcium concentration and a net decrease in AMPA receptor phosphorylation. In contrast, H+VS stimulation evoked larger elevations in calcium concentration, primarily due to the activation of low-voltage activated Ca^2+^ channels upon the release of hyperpolarisation. This led to a substantial increase in AMPA receptor phosphorylation. The role of low-voltage activated Ca^2+^ channels in mediating the high dendritic [Ca^2+^] following periods of hyperpolarisation is in agreement with the finding of Person and Raman [2010] that the induction of hyperpolarisation-gated synaptic plasticity in DCN neurons is dependent on LVCa channel activation.

This hyperpolarisation-gated synaptic plasticity relies on the delicate balance between kinase and phosphatase activity in the post-synaptic dendrite. CaMKII and PP2B compete for activation by calcium-activated calmodulin, and their relevant concentrations are instrumental in determining whether LTP or LTD is induced. This is consistent with previous results [Stefan et al., 2008, Li et al., 2012]. Further downstream, CaMKII and PP1 compete to determine the phosphorylation status both of AMPA receptors and of CaMKII itself. Their bi-stable behaviour determines the bi-stable response of AMPA receptor phosphorylation [Zhabotinsky, 2000, Miller et al., 2005, Pi and Lisman, 2008]). This means that the biochemical reaction system functions as a switch that can produce long-term potentiation or long-term depression in response to changes in calcium concentrations. Our present findings show how the electrophysiological activation of the MVN neuron by patterns of synaptic excitation coupled with inhibition, induces changes in dendritic calcium concentrations and drives the biochemical reaction system to bring about hyperpolarisation-gated plasticity at the vestibular nerve synapse.

The amount and direction of hyperpolarisation-gated synaptic plasticity are dependent in complex ways on the strength and duration of the hyperpolarising stimulus, as well as the relative timing between hyperpolarisation (inhibition) and excitation. Long-term potentiation is observed when hyperpolarisation is sufficiently strong, and when the release from hyperpolarisation coincides with an excitatory stimulus.

Our multi-scale simulation workflow uses the open source tools NEURON [Carnevale and Hines, 2006] and COPASI [Hoops et al., 2006] that work with community-driven standards (NeuroML [Gleeson et al., 2010] and SBML [Hucka et al., 2003]). This provides a useful template for the development of further multi-scale models linking electrophysiological and biochemical models more generally. The modularity of our model means it can also be extended to include more components of the biochemical signalling pathways or a wider neuronal signalling network. It thus has the potential to address a wide range of questions about neural signal integration, post-synaptic biochemical reaction systems and plasticity. While our current model appears to contain the essential components required to express hyperpolarisation-gated synaptic plasticity, further developments are necessary to incorporate additional inputs to MVN neurons that modulate synaptic plasticity, and cellular mechanisms involved in the consolidation of synaptic plasticity through the regulation of gene expression, for example.

## 4 Methods

### 4.1 Biochemical model of signalling pathways underlying synaptic plasticity

The biochemical model used here is based on previous models of calcium signalling and the activation of protein kinases and phosphatases in postsynaptic compartments [Stefan et al., 2008, Li et al., 2012, Mattioni and Le Novère, 2013], starting from Ca^2+^ input and leading to AMPA receptor phosphorylation as a readout. Specifically, we have built on an earlier model by Li et. al [Li et al., 2012], which we helped encode in SBML format, and which is now available on BioModels Database [Li et al., 2010] (BIOMD0000000628). The model follows the basic SBGN reaction scheme introduced in Fig 1B. Accounting for the fact that several of the model components can exist in different functional states [Stefan et al., 2014] and modelling each of those states explicitly, there are a total of 129 molecular species and 678 reactions in the model. Initial concentrations of chemical species were taken from the model by Li et al. [2012]. Concentration of CaMKII was changed to 1 × 10^−5^ M for MVN neurons [Biber et al., 1984]. Besides, we added the reaction: CamR_Ca2_AC + PP2B → CamR_Ca2_AC_PP2B, which is necessary for completeness, but was missing in the model by Li et al. [2012]. We also added PP2B activation to the model which was described in the paper by Li et al. [2012], but was not included in their supplemental model file. For autophosphorylation of CaMKII, we used the polynomial formula used by Li et al. [2012] to compute phosphorylation rates, which accounts for the fact that autophosphorylation proceeds from one active subunit to its neighbour in the CaMKII holoenzyme. However, we slightly modified the rate formula to ensure that the autophosphorylation rate was always greater than 0, in order to be more biochemically accurate. At last, we removed Ca^2+^ buffer proteins from the system, because we already account for calcium buffering when translating between electrical and chemical models (see below).

Initial concentrations of chemical species were taken from previous literature based on rat brains [Biber et al., 1984, Li et al., 2012, Mattioni and Le Novère, 2013] and are presented in table 1. Reaction rates and other parameters were as described by Li et al. [2012].

**Table 1:**
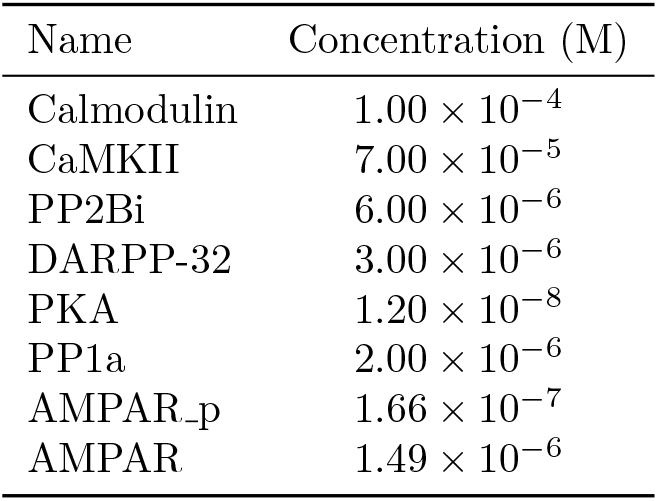
Initial concentration of molecular species in the biochemical model

The biochemical model was edited and run in COPASI 4.16 [Hoops et al., 2006]. It is available in the project’s GitHub repository (https://github.com/YubinXie/multiscale-synaptic-model) in SBML level 2.4 (.xml) format.

### 4.2 Simulation of Chemical model

In order to observe concentration dynamics of key chemical species over time, the Time Course function in COPASI was used. We chose a time interval of 0.0001 s, and the deterministic (LSODA) method. The concentration of Ca^2+^ was either fixed throughout the time course, or altered using the “Events” functionality in COPASI in order to simulate a dynamic calcium signal (see below).

In order to explore equilibrium behaviours at different Ca^2+^ concentrations (Fig. 2), we used the Parameter Scan function in COPASI. Initial Ca^2+^ concentrations ranging from 4 × 10^−10^ M to 4 × 10^−7^ M were scanned, with 151 logarithmic intervals. At each initial Ca concentration, the simulation started from the initial state of the biochemical model and was run for 1600 s with the time interval being 0.0001 s. This was long enough for the system to reach an equilibrium state. The value of Ca^2+^ concentration was fixed at initial value during the whole simulation.

### 4.3 Electrical model of MVN type B neuron

We used a multi-compartmental electrophysiological model of an MVN Type B neuron adapted from Quadroni and Knopfel [1994] and implemented [Graham et al., 2009] in NEURON [Carnevale and Hines, 2006]. The model neuron consisted of a soma, 4 proximal dendrites and 8 distal dendrites, comprising 61 electrical compartments. Each compartment included up to nine active ionic channels (Table 2), and with the exception of the soma also included one excitatory vestibular nerve synapse containing by AMPA receptors and NMDA receptors. The strength of the vestibular nerve synapses was adjusted so that stimulation at a frequency of 100 Hz caused an increase in firing rate of the model MVN neuron similar to that observed experimentally by McElvain et al. [2010].

**Table 2:** Distribution and density of ionic conductances of MVN B neurons[Quadroni and Knopfel, 1994]

| Channels | Maximal conductance per membrane area ($\mu S cm^{-2}$) | | |
| --- | --- | --- | --- |
|  | Soma | Proximal Dendrites | Distal Dendrites |
| Na | 43000 | 2880 | 0 |
| Nap | 23.6 | 38 | 0 |
| H | 66 | 66 | 0 |
| bK(fast) | 37530 | 2572 | 640 |
| AHP | 2716 | 0 | 0 |
| K(slow) | 519 | 406 | 0 |
| A | 755 | 0 | 0 |
| Ca(HVA) | 2385 | 1417 | 350 |
| Ca(LVA) | 166 | 325.5/651/1607* | 50 |
| Na(leak) | 37.8 | 0.7 | 0.71 |
| Ca(leak) | 74.6 | 1 | 1 |
| K(leak) | 166 | 3.69 | 3.68 |

\*The values are for Type B<sub>low\_LVA</sub>, normal Type B, Type B<sub>high\_LVA</sub> MVN neurons, respectively.

We used the two canonical subtypes of Type B MVN neurons modelled by Quadroni and Knopfel [1994] in our simulations. The normal Type B MVN neuron, representing the majority of MVN neurons [Straka et al., 2005], expresses a moderate level of low-voltage activated Ca^2+^ channels (LVA channels, Table 2). By contrast the “Type B_high LV_A” neuron, normally representing some 10% of MVN neurons [Serafin et al., 1991a, Him and Dutia, 2001], expresses a higher level of LVA channels and shows a pronounced low-threshold rebound firing (“low-threshold Ca^2+^ spike”) upon release from hyperpolarisation. While the models of Quadroni and Knopfel [1994] accurately replicate the electrophysiological properties of normal Type B and Type B_high LV_A neurons, it should be noted that the expression of LVA Ca^2+^ channels in MVN neurons and the incidence of post-hyperpolarisation rebound firing is heterogeneous [Serafin et al., 1991a, Straka et al., 2005]. Indeed LVA channel expression in MVN neurons is rapidly upregulated after deafferentation, with the number of Type B_high LV_A neurons increasing significantly during vestibular compensation [Him and Dutia, 2001, Straka et al., 2005]. In this light, to investigate the functional role of LVA channels in the present study, we also modelled a third, hypothetical “Type B_low_LVA_” MVN neuron, where the LVA Ca^2+^ channel density was reduced to 50% of that in normal Type B neurons (Table 2).

### 4.4 Multi-scale interface and stimulation protocols

We applied the stimulation protocols that have been shown experimentally to induce bidirectional plasticity of excitatory synapses in DCN and MVN neurons in vitro [Person and Raman, 2010, McElvain et al., 2010], to determine the evoked dendritic [Ca^2+^] profiles and their effects on synaptic AMPA receptors in the multi-scale model. The vestibular synapses on the proximal and distal dendritic compartments of the MVN model were activated for 550 ms at a frequency of 100 Hz. Repeated periods of vestibular synaptic activation alone (“VS” protocol) have been shown to cause LTD of excitatory synapses in MVN neurons [McElvain et al., 2010]. Alternatively, in the hyperpolarisation + vestibular synaptic activation protocol (H+VS), vestibular synaptic stimulation was paired with hyperpolarisation of the post-synaptic cell for 250 ms, by the injection of inhibitory current into all of the dendritic compartments. This pattern of stimulation was shown to induce LTP of vestibular nerve synapses in MVN neurons [McElvain et al., 2010].

Values of [Ca^2+^] in the mid-proximal dendrites during the normal resting activity of the MVN neuron and in response to the VS and H+VS stimulation protocols were obtained from NEURON, and used as the input to the biochemical model in COPASI. The [Ca^2+^] values obtained from NEURON were scaled down by a factor of 50, to account for Ca buffering and sequestration processes not explicitly included in our biochemical model (Fig.11).

**Figure 11:**
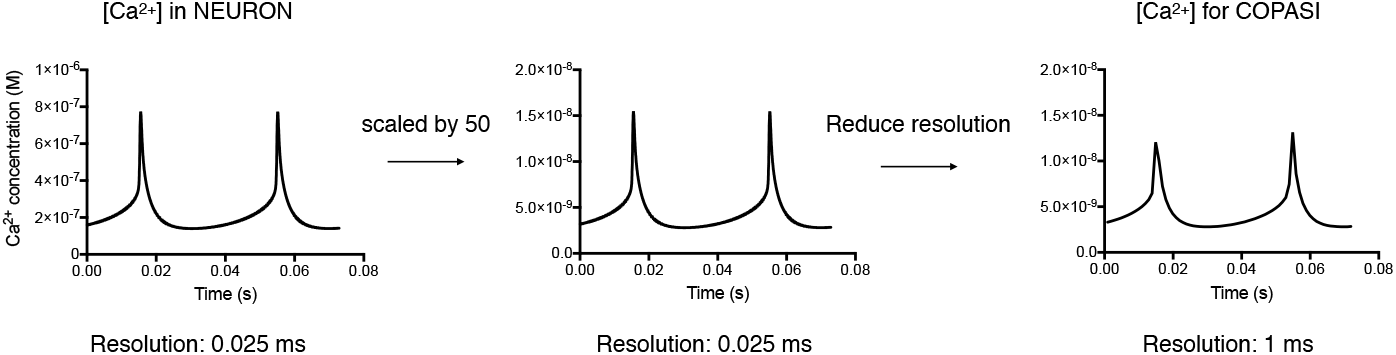
Demonstration of [Ca^2+^] conversion from electrical modelling tool NEURON to biochemistry modelling tool COPASI. Firstly, the [Ca^2+^] values obtained from NEURON were scaled down by a factor of 50, to account for Ca^2+^ buffering and sequestration processes not explicitly included in our biochemical model. Then, the time resolution was scaled up by a factor of 40, given the relatively slower dynamics in biochemistry system when compared to the electrical system.

**Figure 12:**
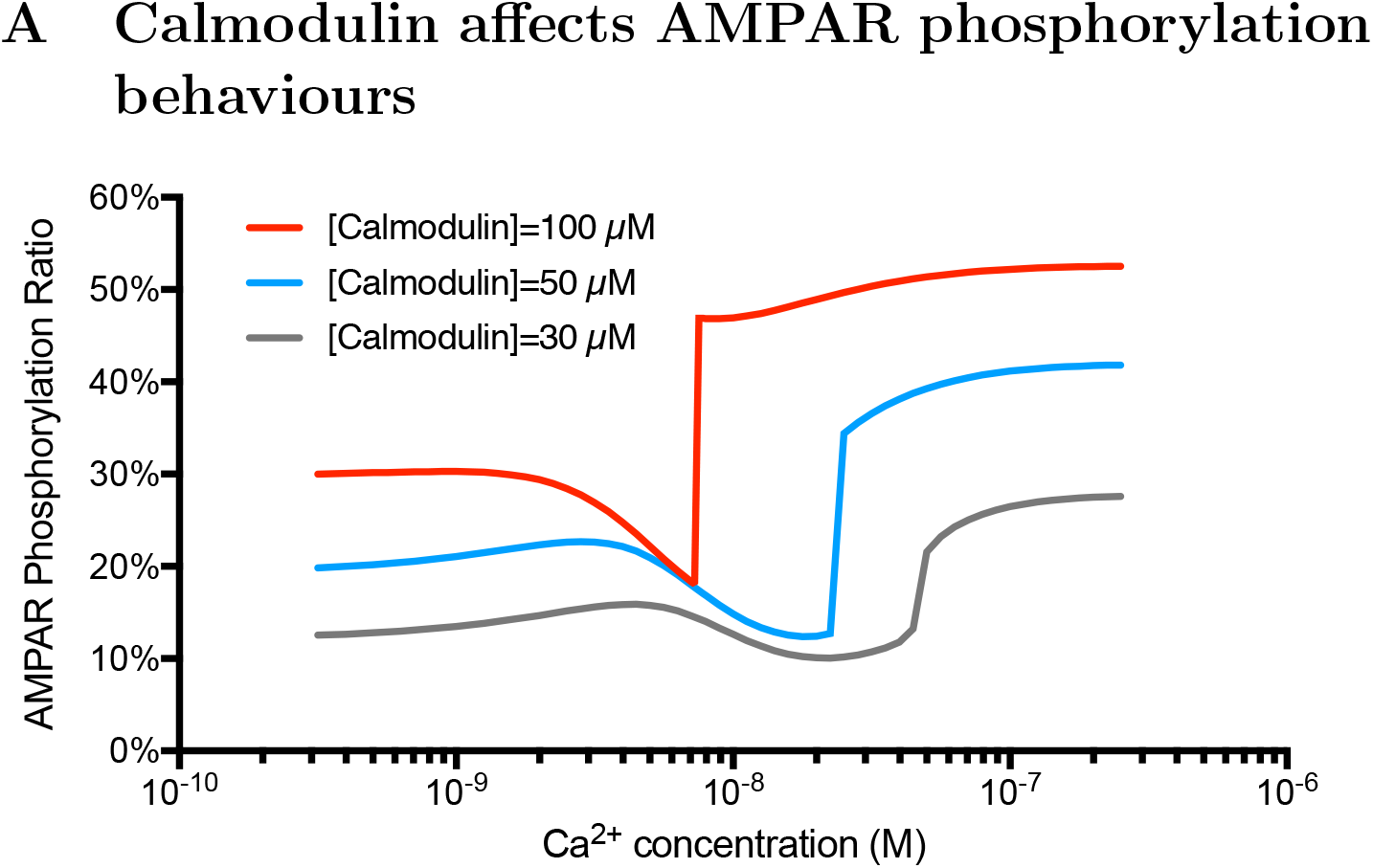
The relationship between AMPAR phosphorylation ratio and Ca^2+^ concentration when calmodulin being 100 μM, 50 μM, 30 μM. A higher concentration of calmodulin gives a faster and stronger switch-like behaviour of AMPAR phosphorylation when the calcium concentration increases.

This scaling factor brought the [Ca^2+^] values calculated in NEURON into the physiological range of intracellular Ca^2+^ concentration known from experimental studies [Sharma et al., 1995]. In addition, the chemical model has a much lower time resolution than the electrical model, with a simulation time step of 1 ms vs. 0.025 ms in the electrical model. We therefore calculated the average of 40 successive values of [Ca^2+^] in the NEURON model, to provide the COPASI model with an input value of [Ca^2+^] at every 1 ms time step (Fig.11). After [Ca^2+^] scaling and time resolution transformation, the resulting [Ca^2+^] values was added to the COPASI input file as an “Event”. Then, the biochemical model could be run in COPASI. All the process was automatically done by a Python script (provided here: https://github.com/YubinXie/multiscale-synaptic-model). One thing to note is that, before adding the stimulation events into COPASI model, the COPASI model should run for 1000 s with the Ca^2+^ being the average Ca^2+^ of the corresponding (Type B, Type B_;Low_LVA_, Type B_high_LVA_) resting neurons.

#### Regular H + VS simulation protocol (Fig. 3, 5, 4, 6)

In a 1000 ms window, 100Hz 550 ms VS stimuli starts at 250 ms and ends at 800 ms. 475 pA inhibitory stimuli (hyperpolarisation) starts at 250 ms and ends at 500 ms. This 1000 ms simulation is constantly repeated until the simulation ends (360 s). Inhibitory stimuli of 475 pA on Type B_high LV_A neurons gave a huge [Ca^2+^] rebound, which is too huge to be visualized in a reader-friendly way. In Fig. 5, 6, 320 pA inhibitory stimuli was used in Type B_high LVA_ neurons.

#### Hyperpolarisation strength simulation protocol (Fig. 8)

In a 1000 ms window, a 100 Hz 550 ms VS stimulus starts at 250 ms and ends at 800 ms. Inhibitory stimulus (hyperpolarisation) with strength from 0 pA to 600 pA starts at 250 ms and ends at 500 ms. This 1000 ms window is constantly repeated until the simulation ends (600 s).

#### Hyperpolarisation duration simulation protocol (Fig. 9)

In a 1000 ms window, 100 Hz, a 550 ms VS stimulus starts at 250 ms and ends at 800 ms. A 475 pA inhibitory stimulus (hyperpolarisation) with duration from 0ms to 550 ms starts at the same time. This 1000 ms window is constantly repeated until the simulation ends (600 s).

#### Hyperpolarisation timing simulation protocol (Fig. 10)

To allow a large range of timing scanning, here we used a 1500 ms time window. In this window, a 100 Hz 250 ms VS stimulus starts at 250 ms and ends at 500 ms. A 250 ms inhibitory stimulus (hyperpolarisation) with strength 475 pA starts ranging form 500 ms earlier to 250 ms later than the VS stimulus. This 1500 ms simulation is constantly repeated until the simulation ends (600 s).

Individual simulations were run on standard desktop and laptop computers. Sets of simulations for figures 8 to 10 were run on the Eddie Compute Cluster at the University of Edinburgh.

### 4.5 Reproducibility

All the models and necessary scripts that were used in this paper can be found in the following GitHub repository: https://github.com/YubinXie/multiscale-synaptic-model. Detailed guides are provided to reproduce all the figures.

## 5 Acknowledgements

The authors thank Nicolas Le Novère for discussions on the System Biology Graphical Notation form of the biochemical model schematic. We thank Lu Li, Pinar Pir, and Varun B. Kothamachu for discussion and input on the System Biology Markup Language version of the biochemical model. We also thank Sven Sahle for discussions on the COPASI software. Gary Westbrook provided helpful feedback on an earlier version of this manuscript. Marcel Kazmierczyk’s work on this project was supported by a vacation scholarship from Medical Research Scotland.

